# Association between schizophrenia polygenic score and psychotic symptoms in Alzheimer’s disease: meta-analysis of 11 cohort studies

**DOI:** 10.1101/528802

**Authors:** Byron Creese, Evangelos Vassos, Sverre Bergh, Lavinia Athanasiu, Iskandar Johar, Arvid Rongve, Ingrid Tøndel Medbøen, Miguel Vasconcelos Da Silva, Eivind Aakhus, Fred Andersen, Francesco Bettella, Anne Braekhus, Srdjan Djurovic, Giulia Paroni, Petroula Proitsi, Ingvild Saltvedt, Davide Seripa, Eystein Stordal, Tormod Fladby, Dag Aarsland, Ole A. Andreassen, Clive Ballard, Geir Selbaek, on behalf of the AddNeuroMed consortium and the Alzheimer’s Disease Neuroimaging Initiative

## Abstract

**Background:** Psychosis (delusions and hallucinations) is common in Alzheimer’s disease (AD) and associated with worse clinical outcomes including accelerated cognitive decline and shorter time to nursing home admission. Atypical antipsychotics have limited efficacy which, along with emerging genomic research, suggests some overlapping mechanisms with other disorders characterized by psychosis, like schizophrenia. In this study, we tested whether polygenic risk score (PRS) for schizophrenia was associated with psychotic symptoms in AD.

**Methods:** Schizophrenia PRS was calculated using Psychiatric Genomics Consortium data at 10 GWAS p-value thresholds (*P*_*T*_) in 3,173 AD cases from 11 cohort studies. Association between PRS and AD psychosis status was tested by logistic regression in each cohort individually and the results meta-analyzed.

**Results:** The schizophrenia PRS was associated with psychosis in AD at an optimum *P*_*T*_ of The strongest association was for delusions where a one standard deviation increase in PRS was associated with a 1.17-fold increased risk (95% CI: 1.07-1.3; p=0.001).

**Conclusion:** These new findings point towards psychosis in AD – and particularly delusions – sharing some genetic liability with schizophrenia, and support a transdiagnostic view of psychotic symptoms across the lifespan.

## Introduction

Psychosis in Alzheimer’s disease (AD-P) - broadly comprising delusions and hallucinations - is experienced by up to 50% of people over the course of the illness (1). AD-P is associated with accelerated cognitive decline (independent of disease duration), higher mortality rates and distress to both the people with dementia and their carers (2-4). Moreover, there are wider societal implications with long-term follow up studies indicating that AD-P is associated with a shorter time to nursing home care (5). Despite these compelling reasons for effective management, there is a critical treatment gap, with no licensed treatments available in many jurisdictions. Atypical antipsychotics – developed for other conditions characterized by psychosis – are frequently used (in many countries off label) but have limited benefits and are associated with considerable harms, including a 1.5-to 1.8-fold increase in mortality and a 3-fold increase in stroke (6).

There are only two new antipsychotic compounds in phase II or later stages of development (pimavanserin and MP-101) but both are refinements of existing mechanisms of action of atypical antipsychotics or other compounds targeting mechanisms relevant to schizophrenia (e.g. 5HT2A, mGluR2/3) (7). The limited understanding of the biological mechanisms underpinning AD-P represents a major challenge to the effective targeting of existing treatments and the development of novel compounds.

One key question is whether some or all of the psychotic symptoms experienced by people with AD-P have a similar basis to the psychotic symptoms experienced by people with schizophrenia. Phenomenologically the symptoms are different; in AD-P visual hallucinations are more common than auditory hallucinations, delusions are usually simple, and first rank symptoms of schizophrenia are very rare. Despite the different phenomenology, atypical antipsychotics confer some treatment benefits in some cases of AD-P (8), suggesting at least some potential overlap.

This transdiagnostic hypothesis, proposing a mechanistic overlap between AD-P and schizophrenia, is gaining some traction (9). Given the high heritability of schizophrenia (10) and of AD-P (estimated at 81% and 61% respectively) (11), it is logical to look for common genetic underpinnings of the two disorders. In a small study, a copy number variant (CNV) with significant overlap of a duplicated region implicated in schizophrenia and autism (16p11.2) was found in two of 440 AD-P cases but not in AD without psychosis or in those with more occasional symptoms (12). Linkage studies have also implicated regions of the genome in AD-P that have been identified in schizophrenia (13, 14). Another approach is to examine whether polygenic risk for schizophrenia is associated with AD-P. Work in this area is limited to only one recent study which reported that a genetic risk score comprising 94 SNPs reaching genome wide significance for association with schizophrenia was lower in AD-P compared with AD without psychosis (15). While this study represents an important step in AD-P research, a full genome-wide polygenic risk score (PRS) approach is warranted (16, 17).

Another largely unexplored avenue in AD-P genetic research relates to the split of delusions and hallucinations. Although the two symptoms frequently co-occur in AD, there is evidence from longitudinal cohort studies indicating that 10-20% of people experience hallucinations without delusions and that the two symptoms are associated with different clinical outcomes (2, 18), suggesting the presence of two distinct clinical phenotypes. While it is commonplace to separate out composite psychotic symptoms in neuroimaging studies of AD-P (19, 20), their separate genetic associations have not yet been examined in any large-scale AD studies leveraging GWAS data (21). This is a particularly relevant issue when assessing genetic overlap with schizophrenia where the emerging evidence from neuroimaging and the clinical similarity supports the hypothesis that shared etiology would be specific to delusions.

We conducted an analysis of the relationship between genetic liability for schizophrenia and psychotic symptoms in AD with two principal objectives; firstly, we tested whether a polygenic risk score (PRS) for schizophrenia was associated with psychosis in AD and secondly, we examined whether refining the psychosis phenotype to focus on delusions would lead to a stronger association.

## Materials

### Cohorts

AD-P target data consisted of 3,173 AD cases from 11 cohort studies in Europe and the USA: AddNeuroMed (22) (Europe), Alzheimer’s Disease Neuroimaging Initiative (23) (ADNI; USA), Istituto di Ricovero e Cura a Carattere Scientifico (IRCCS 1; Italy), Health and Memory Study in Nord-Trøndelag (24) (HMS; Norway), Resource Use and Disease Couse in Dementia (25) (REDIC; Norway), Norwegian registry of persons assessed for cognitive symptoms (26) (NorCog; Norway), Samhandling mellom avdeling for alderspsykiatri og kommunale sykehjem (SAM-AKS; Norway), The Dementia Study in Northern Norway (27) (NordNorge, Norway), Progression of Alzheimer’s Disease and Resource Use (28) (PADR; Norway), The Dementia Study in Western Norway (29) (DemVest; Norway); and data from the National Alzheimer’s Disease Coordinating Center (NACC) and the National Institute on Aging Genetics Data Storage Site (NIAGADS), Table 1). Full cohort details are contained in the supplementary material and the Norwegian cohorts are also described in the latest GWAS of Alzheimer’s disease (30).

**Table 1:** Baseline demographics by cohort

|  | N | Age |  | Gender<br>% male | MMSE |  | Assesment<br>scale | Array |
| --- | --- | --- | --- | --- | --- | --- | --- | --- |
|  |  | Mean | SD |  | Mean | SD |  |  |
| AddNeuroMed | 224 | 77 | 7.2 | 35 | 20 | 4.7 | NPI | Illumina 610 |
| ADNI | 248 | 76 | 7.2 | 59 | 24 | 2.5 | NPI-Q | Illumina OmniExpress |
| DemVest | 80 | 76 | 6.6 | 33 | 24 | 2.4 | NPI | Illumina OmniExpress |
| IRCCS 1 | 326 | 79 | 7.1 | 41 | 12 | 6.4 | NPI | Illumina GSA |
| HMS | 178 | 87 | 6.0 | 26 | 13 | 6.5 | NPI | Illumina OmniExpress |
| NorCog | 623 | 75 | 9.0 | 43 | 22 | 4.3 | NPI-Q | Illumina OmniExpress |
| NordNorge | 133 | 80 | 6.7 | 41 | 23 | 4.4 | NPI | Illumina OmniExpress |
| PADR | 106 | 76 | 6.6 | 33 | 21 | 4.3 | NPI-Q | Illumina OmniExpress |
| REDIC | 327 | 85 | 7.0 | 34 | 16 | 6.4 | NPI | Illumina OmniExpress |
| SAM-AKS | 92 | 86 | 6.4 | 28 | 16 | 5.0 | NPI | Illumina OmniExpress |
| NACC | 836 | 78 | 8.2 | 50 | 20 | 7.1 | NPI-Q | Illumina 660/Omni<br>Express |
| **TOTAL** | **3173** | **78** | **8.6** | **42** | **19** | **6.7** | **-** | **-** |

NPI: Neuropsychiatric Inventory (full version); NPI-Q: Neuropsychiatric Inventory- Questionnaire; MMSE: Mini Mental State Examination

Some data used in the preparation of this article were obtained from the Alzheimer’s Disease Neuroimaging Initiative (ADNI) database (adni.loni.usc.edu). The ADNI was launched in 2003 as a public-private partnership, led by Principal Investigator Michael W. Weiner, MD. The primary goal of ADNI has been to test whether serial magnetic resonance imaging (MRI), positron emission tomography (PET), other biological markers, and clinical and neuropsychological assessment can be combined to measure the progression of mild cognitive impairment (MCI) and early Alzheimer’s disease (AD). For up-to-date information, see www.adni-info.org.

### AD clinical assessments

Diagnosis of AD was performed according to ICD-10 etiological diagnosis, NINCDS-ADRDA criteria or clinical diagnosis by psychiatrist or geriatrician. Longitudinal data was available for 6 cohorts (ADNI, AddNeuroMed, DemVest, NordNorge, PADR, REDIC, NACC) and psychotic symptom classification was on the maximum amount of follow up data available. Any cases with a history of bipolar disorder or schizophrenia were excluded. For NorCog, PADR, REDIC, SAM-AKS, NACC and ADNI the necessary information on psychiatric history was extracted from source study data resulting in 3, 1, 2, 1, 31 and 1 exclusions respectively. For AddNeuroMed, DemVest, IRCCS 1 and NordNorge this was an exclusion criteria applied at entry to those individual studies. No information about psychiatric history was available for the HMS study. Dementia severity was assessed in all cohorts by Mini Mental State Examination (MMSE) and psychotic symptoms were assessed by the Neuropsychiatric Inventory (NPI) or its short version, the Neuropsychiatric Inventory Questionnaire (NPI-Q), two widely used validated instruments (31). Psychotic symptoms are rated on the basis of items A (delusions) and B (hallucinations) of the NPI and NPI-Q. In the full NPI, neuropsychiatric symptoms are coded as present or absent first. If rated present they are further scored according to their frequency (1-4) and severity (1-3) with the resulting scores multiplied to give an overall rating (i.e. possible scores are 1,2,3,4,6,8,9 and 12 with 0 indicating no symptoms). The NPI-Q is rated only on a scale of 0 to 3 according to the severity of the symptom. Both scales have been designed to be completed by verbal interview with a proxy informant who knows the person with AD well. Several diagnostic criteria for AD psychosis have been proposed but none have been adopted clinically, meaning that where in other psychiatric disorder medical records can be screened in AD-P this would be unreliable and ratings on specific validated assessment scales must be used. We thus undertook examination of three related but progressively more homogenous psychotic phenotypes:

1. *Psychosis wide:* Psychosis present: the presence of delusions or hallucinations (>0) at any point; No psychosis: no evidence of delusions or hallucinations at any point in follow up.
2. *Psychosis narrow:* Psychosis present: the presence of delusions or hallucinations (>0) at any point during follow up; No psychosis: here, an additional level of screening was applied to those rated as having no delusions or hallucinations. In these cases, if an individual was psychosis-free and had not yet reached the moderate to severe stages of dementia (defined as MMSE<20 at their last assessment) they were excluded from the analysis. This is a similar approach to that used in most previous AD-P genetic research (15, 32).
3. *Delusions narrow:* Delusions present: the presence of delusions (>0) at any point during follow up. Thus, the delusion group was the psychosis group above with any cases with hallucinations only removed. No delusions: as per psychosis narrow. We did not analyze a hallucinations group as most people with hallucinations also experience delusions so any grouping on this basis would overlap substantially with the delusions group.

### Genotyping and QC

The genotyping chips used are detailed in Table 1. Raw genotype data for individual cohorts underwent appropriate QC steps (implemented in PLINK). SNPs with a minor allele frequency ≤5% and a Hardy Weinberg equilibrium p < 10^-5^ were excluded. The SNP and individual genotype failure threshold was set at 5% and individuals with mean heterozygosity ±3 standard deviations were excluded. The analysis was restricted to individuals of European ancestry using genetic principal components computed by EIGENSTRAT. Related (pi-hat >0.2) or duplicate individuals both within and between cohorts were excluded. Phasing (EAGLE2) and imputation (PBWT) was done via the Sanger Imputation Service using the Haplotype Reference Consortium (r1.1) reference panel on all cohorts. After imputation only SNPs with an imputation quality (INFO) score >0.4 and MAF >0.05 were retained. 4,895,913 SNPs common across all ten cohorts were retained to compute polygenic risk scores PRS.

The most recently published schizophrenia GWAS data from the Psychiatric Genomics Consortium was used as base data to generate PRS in the target AD sample (17). SNPs with MAF<0.1, INFO<0.9 and indels were excluded from the base dataset to leave only the most informative SNPs and only one SNP from the extended MHC region was included (33).

### Analysis

PRS were generated in PRSice (34) at the following 10 GWAS p-value thresholds (*P*_*T*_): 5×10^−8^, 1×10^−5^, 1×10^−4^, 1×10^−3^, 0.01, 0.05, 0.1, 0.2, 0.5 and 1. Clumping was performed (250kb, r2>0.1) to retain only the SNP with the strongest association in each window. The resulting PRS were standardized for the analysis.

All statistical analysis was implemented in R. For each cohort 11 logistic regression models (one per *P*_*T*_) were run with each of the previously defined psychosis phenotypes as the binary outcome and the first 10 ancestry principal components included as covariates. Logistic regression assumptions were confirmed using the R ‘car’ package. Proportion of variance explained (*R*^*2*^) by PRS was determined by subtracting the Nagelkerke’s pseudo-*R*^*2*^ of the null model from that of the full model. Regression coefficients for each *P*_*T*_ across all cohorts were then included in random effects meta-analyses to account for between-study variation in data collection protocols, frequency of psychosis and dementia severity. Meta-analysis was undertaken using the ‘rma’ function in the ‘metafor’ package using the REML method (35).

## Results

On average across all eleven cohorts, individuals were in the mild-moderate stages of dementia at first assessment (mean MMSE of 19). Mean MMSE by cohort ranged from an MMSE of 12 (IRCCS 1) to 24 (ADNI) and this was a correlate of the prevalence of psychosis in each cohort (note the denominator would be the overall cohort N in Table 1), with cohorts that contained individuals with more severe dementia typically having a higher proportion of people with psychosis. Between cohorts, mean age at baseline ranged from 75 to 87 years and the proportion of male participants ranged from 26% to 59%.

Frequency of the three phenotypes investigated by cohort is shown in Table 2. Of the 3,3173 individuals screened, 1,129 (36%) had psychosis (wide definition group). Of the 2,044 who were rated as having no psychosis based on their assessment scale result alone, 913 had not yet reached the moderate stages of disease and so were excluded, thus forming the psychosis narrow group. 1,129 psychosis cases and 1,131 AD no psychosis ‘controls’ were included in the analysis of the narrow phenotype of psychosis. 951 cases met the criteria for having delusions narrow.

**Table 2:**
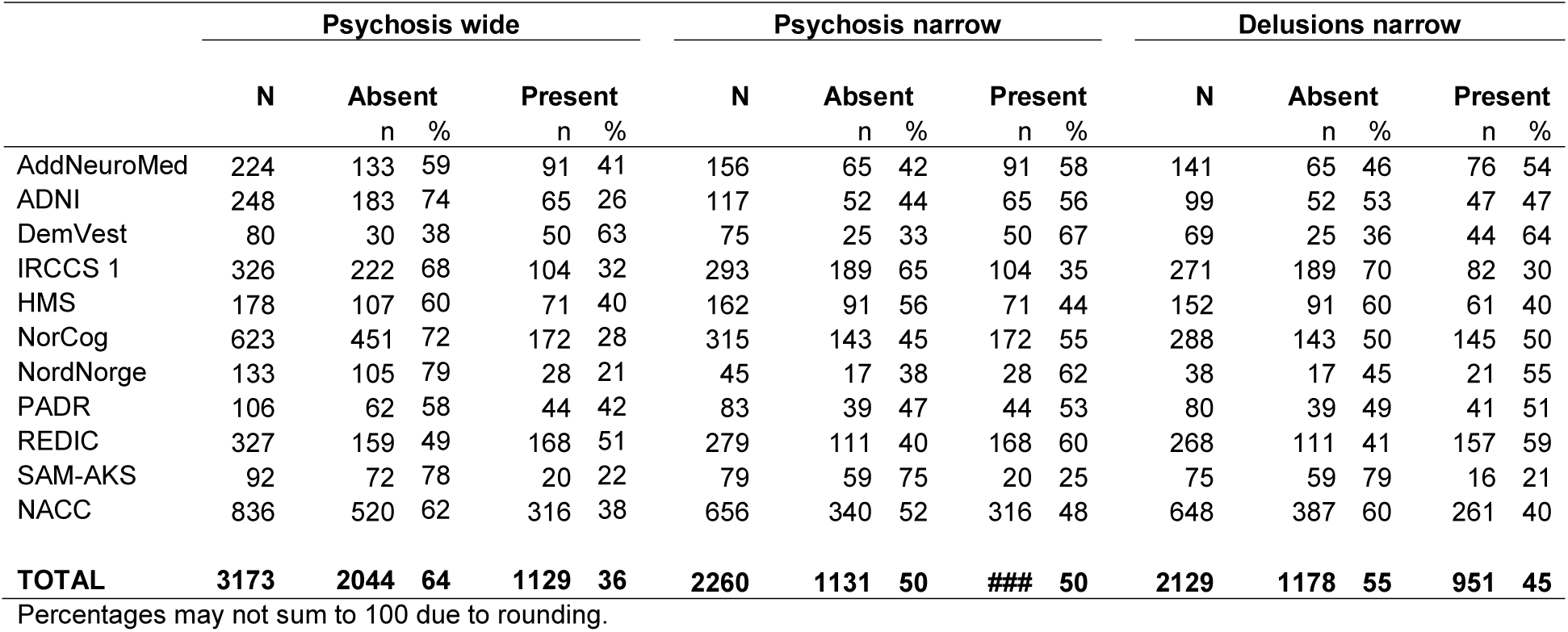
Frequencies of symptoms by cohort for the three psychosis phenotypes

### Schizophrenia PRS is associated with AD psychosis status

After clumping, 76,213 independent variants were available for computing PRS. Random effects meta-analysis across the 11 cohorts showed schizophrenia PRS at *P*_*T*_=0.01 was significantly associated with symptom status with effect size estimates increasing progressively across psychosis wide, psychosis narrow and delusions narrow (1.15 95% CI:1.04-1.27, p=0.005 1.19 95% CI:1.06-1.34 p=0.004,1.2 95% CI:1.06-1.36, p=0.003 respectively), see Figure 1 and Table 3. PRS was significantly associated with both the psychosis narrow and delusions narrow phenotypes at every *P*_*T*_ >0.01. The largest effect size was observed in the delusions narrow group. Overall, there was no evidence of significant heterogeneity as indicated by *I*^*2*^ statistics, low heterogeneity was present at P_T_≥0.5 in the psychosis wide phenotype.

**Figure 1:**
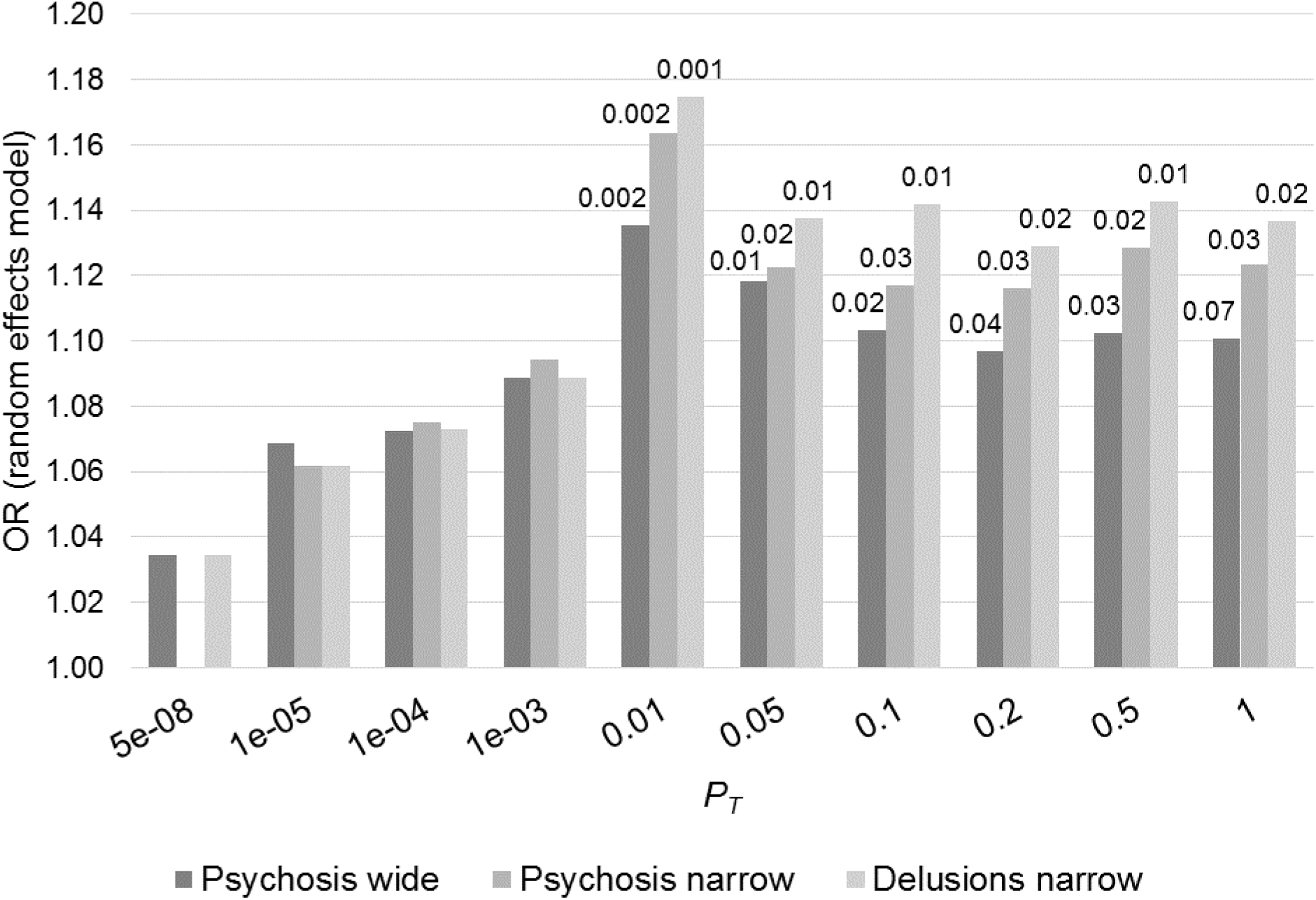
Odds ratios from random effects meta-analysis of AD psychosis wide, narrow and delusions narrow association with schizophrenia PRS. Each bar represents PRS composed of markers at 10 different schizophrenia GWAS p-value thresholds (*P*_*T*_). P-values shown above each bar

**Table 3:** Random effects meta-analysis results for psychosis wide definition across the 10 schizophrenia GWAS thresholds.

| **P<sub>T</sub>** | **nSNPs** | **Psychosis wide** |  |  |  |  | **Psychosis narrow** |  |  |  |  | **Delusions narrow** |  |  |  |  |
| --- | --- | --- | --- | --- | --- | --- | --- | --- | --- | --- | --- | --- | --- | --- | --- | --- |
|  |  | **OR** | **95% CI** |  |  | **P** | **OR** | **95% CI** |  |  | **P** | **OR** | **95% CI** |  |  | **P** |
| 5x10<sup>-08</sup> | 125 | 1.03 | 0.96 | - | 1.12 | 0.40 | 1.00 | 0.91 | - | 1.09 | 0.99 | 1.03 | 0.94 | - | 1.14 | 0.48 |
| 1x10<sup>-05</sup> | 511 | 1.07 | 0.98 | - | 1.17 | 0.13 | 1.06 | 0.97 | - | 1.16 | 0.20 | 1.06 | 0.97 | - | 1.17 | 0.21 |
| 1x10<sup>-04</sup> | 1147 | 1.07 | 0.97 | - | 1.19 | 0.19 | 1.08 | 0.97 | - | 1.20 | 0.18 | 1.07 | 0.97 | - | 1.18 | 0.17 |
| 1x10<sup>-03</sup> | 2922 | 1.09 | 0.98 | - | 1.21 | 0.11 | 1.09 | 0.98 | - | 1.22 | 0.10 | 1.09 | 0.99 | - | 1.20 | 0.09 |
| 0.01 | 8709 | 1.14 | 1.05 | - | 1.23 | 0.002 | 1.16 | 1.06 | - | 1.28 | 0.002 | 1.17 | 1.07 | - | 1.30 | 0.001 |
| 0.05 | 19656 | 1.12 | 1.03 | - | 1.22 | 0.01 | 1.12 | 1.02 | - | 1.24 | 0.02 | 1.14 | 1.03 | - | 1.26 | 0.01 |
| 0.1 | 28143 | 1.10 | 1.01 | - | 1.20 | 0.02 | 1.12 | 1.01 | - | 1.23 | 0.03 | 1.14 | 1.03 | - | 1.27 | 0.01 |
| 0.2 | 40253 | 1.10 | 1.01 | - | 1.20 | 0.04 | 1.12 | 1.01 | - | 1.23 | 0.03 | 1.13 | 1.02 | - | 1.25 | 0.02 |
| 0.5 | 61727 | 1.10 | 1.01 | - | 1.21 | 0.03 | 1.13 | 1.02 | - | 1.25 | 0.02 | 1.14 | 1.03 | - | 1.27 | 0.01 |
| 1 | 76213 | 1.10 | 0.99 | - | 1.22 | 0.07 | 1.12 | 1.01 | - | 1.25 | 0.03 | 1.14 | 1.02 | - | 1.26 | 0.02 |

OR: Odds ratio; odds ratio estimates may differ slightly from those represented in Figure 1 due to rounding

In the individual cohort analysis, we observed that in most cases schizophrenia PRS was higher in cases with psychotic symptoms (all three definitions) than those without; albeit not significantly, possibly due to the small sample sizes (Supplementary material). A forest plot of individual study estimates for delusions narrow at *P*_*T*_=0.01, the strongest association found in the above meta-analysis, is shown in Figure 2. A similar plot at *P*_*T*_=1 for comparison is shown in the Supplementary material along with plots for psychosis wide and psychosis narrow phenotypes. The effect estimates of nine of the 11 studies were in the same direction (OR>1) (Figure 2). The highest *R*^*2*^ estimate was 2.9% (AddNeuroMed) and the lowest was <0.1% (IRCCS 1). An overall *R*^*2*^ of 0.08% was estimated by calculating the weighted average *R*^*2*^ across the 11 studies.

**Figure 2:**
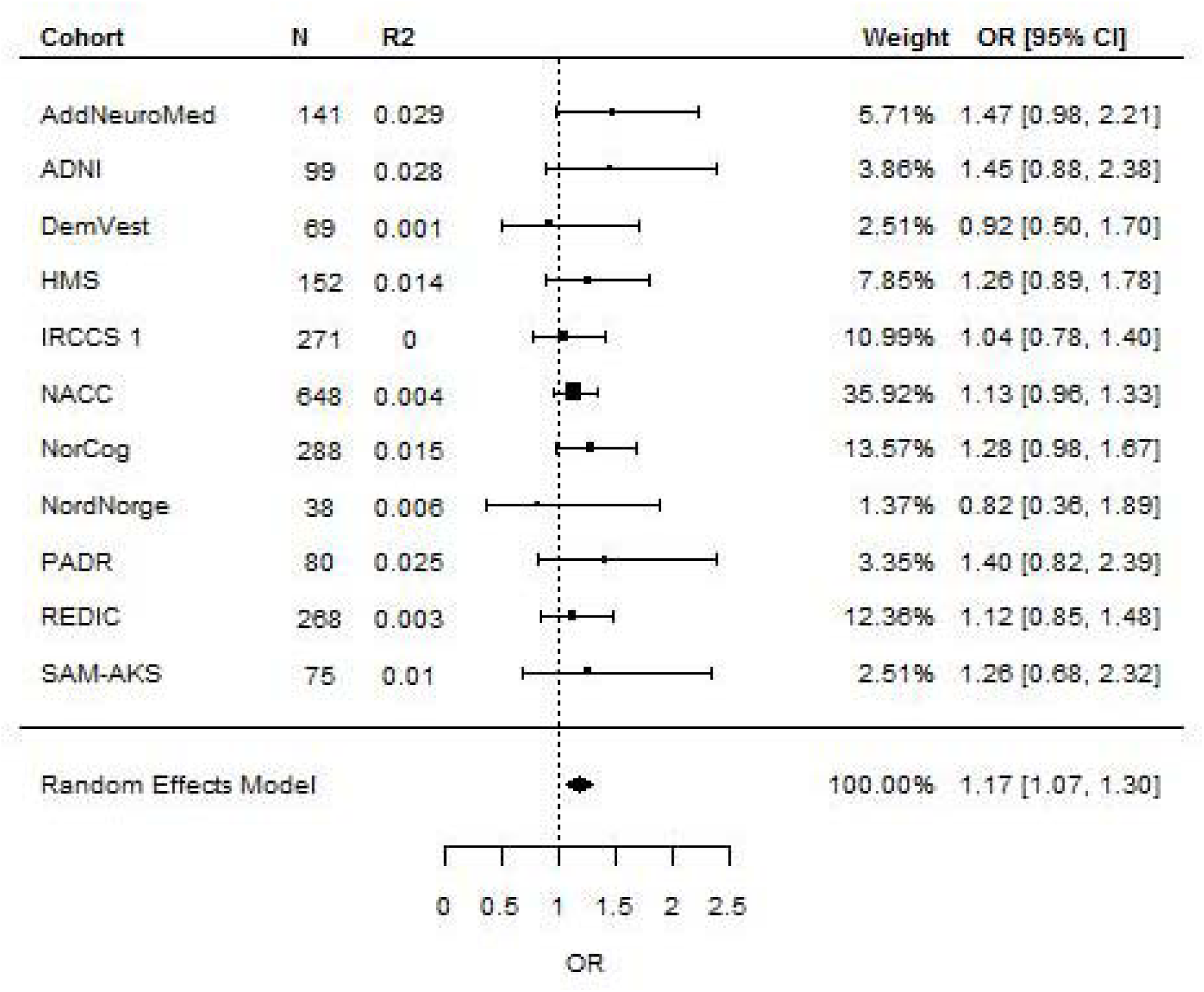
Forest plot of meta-analysis of delusions narrow for PRS calculated at PT=0.01 (i.e. 8,709 SNPs). Overall estimate from random effects model is represented by the diamond below the individual study estimates.

## Discussion

We set out to examine whether genetic risk for AD-P is attributable to common schizophrenia variants. Using polygenic scoring we found that schizophrenia PRS was associated with AD-P in a collection of over 3,000 well characterized cases and that there were some differences in effect size according to the phenotypes examined. The largest effect size was observed at *P*_*T*_=0.01 which was associated with a 1.14, 1.16 and 1.17-fold (per standard deviation increase in PRS) increased risk of psychosis (wide), psychosis (narrow) and delusions (narrow) respectively. The larger effect sizes for the narrow symptom definition phenotypes were observed despite a reduction in the number of cases analyzed. In all, these new findings suggest that psychotic symptoms in AD are part of a spectrum of related psychotic syndromes across the lifespan. Although the overlap between these two phenotypes is only modest, with the *R*^*2*^ estimates being less than 1%, this should be seen in the context of the schizophrenia PRS explaining around 2.5% of the variance in bipolar disorder and 1% in MDD in the last cross-disorder analysis of the Psychiatric Genomics Consortium (36).

Three possible conclusions could be drawn from the finding that the strongest association was observed in the delusions group (i.e. when we removed cases with hallucinations only). Firstly, this may point towards a subset of AD-P patients that have a more schizophrenia-like phenotype. More work is needed to investigate whether further diagnostic refinements to AD-P syndrome definitions are necessary, which may provide a more robust approach for pharmacological intervention trials. Related to this, from a methodological point of view, we show that there is a need for future studies in AD to consider delusions and hallucinations separately. We cannot say from this study that there is no genetic association between hallucinations in AD and schizophrenia in these cohorts but the evidence at present suggests a weaker association than for delusions. One might speculate that this is due to visual hallucinations in AD being more often the result of a broader range of causes (e.g. visual hallucinations due to medication or delirium) than delusions, thus introducing more noise into the phenotype. The final wider implication is related to the schizophrenia PRS being associated with a broad spectrum of psychotic disorders and personality traits (36-38). Our findings extend this research and support a transdiagnostic explanation of delusions, which reaches into neurodegenerative disease and is underpinned by a degree of common genetic liability.

A key strength of our study is the detailed phenotyping with longitudinal data being available in six of the ten cohorts. Phenotyping of AD-P is challenging but the size of our pooled sample and the availability of longitudinal data in half of the cohorts increases our confidence in a robust phenotype. In the absence of any universally used clinical or research diagnostic criteria for psychosis - which would make medical record screening feasible - we followed previous research by taking extra measures to screen the ‘control’ groups. This created two additional phenotypes which removed any cases in the mild stages of disease who had not yet developed symptoms (the cumulative prevalence of psychosis in AD is around 50% (1), peaking in the moderate-severe stages of the disease). This approach has been used in most previous genetic research but our extension to focus on delusions is novel, and, that further refinements to the phenotype strengthened the association is consistent with genetic studies of other polygenic traits (39). For one study (HMS) data on history of major psychiatric conditions were not available. It is possible that some individuals with schizophrenia were present in this cohort however HMS is a nursing home-based cohort with a mean age of 87. Therefore, given that schizophrenia has an overall population prevalence of 1% and a life expectancy is 15-20 years less than the general population, it is highly unlikely that the number would be more than one or two out of 178 people in the HMS cohort (this is also supported by the very small numbers we found among the other studies we screened). With 3,173 samples, this is, to our knowledge, the largest analysis of AD-P to exploit GWAS data (32). We acknowledge that using different cohorts has led to some variability due to sampling in the phenotype examined however in all cases symptoms were assessed by one of two standardized tools (either the NPI or NPI-Q). These are two different versions of the same scale, which are strongly correlated and have good between-rater and test-retest reliability, particularly for the psychosis items (31, 40). It is important to acknowledge that there are no single cohorts which are large enough to conduct an analysis of this kind and because of sampling and protocol variations across the individual studies we ensured an appropriate analysis was implemented to account for this variability. We had access to raw individual-level clinical and genotype data, allowing us to run the same regression models in each study. This included undertaking the same QC across cohorts, imputing all chip data to the same reference panel and analyzing only SNPs present across all cohorts. After ensuring this standard process was followed for each cohort we ran a random effects meta-analysis, allowing for the effect of the PRS on psychotic symptoms to vary across studies. In all, and in the absence of a single large enough study, we believe these measures provide the most robust alternative.

It is important to note that our findings are in contrast with a previous study which examined a more genetic risk score at a more conservative P_T_, comprised of 94 genome-wide significant schizophrenia SNPs (15). We were unable to replicate this association in any of our cohorts, which were independent of this previous study, or in the overall meta-analysis. Our study is a similar size to both the discovery and replication cohorts in this previous study, and the NACC data was used in both. Given that a PRS with only 94 SNPs will be a less powerful predictor than a full genome-wide score it is possible larger studies will be needed to confirm associations at this more conservative *P*_*T*_. Nevertheless, schizophrenia is highly polygenic; tens of thousands of markers explain only 18% of the variance in case-control status and 7% of the variance on the liability scale, while for optimum cross-trait case-control (e.g. schizophrenia and bipolar) prediction many thousands more SNPs are required [19, 20]. Accordingly, it is likely that a full account of association between schizophrenia and AD-P should exploit the full polygenic nature of schizophrenia. Our study is the first to do this and the findings represent an important further step towards a complete account of the relationship between common schizophrenia variants and AD-P. Another important milestone will be an appropriately powered discovery GWAS of AD-P. All of these points underscore the need for increasing samples sizes in this field, and provide a strong motivation for continued efforts towards this goal. To this end we have included individual regression results for our analysis in the supplementary material.

In summary, these findings support common genetic underpinnings between schizophrenia and delusions in AD, suggesting common etiopathogenic mechanisms. This provides a strong rationale for further work to build a clearer clinical and biological understanding of the psychosis syndrome in AD, an urgently needed step for better management and treatment development.

## Acknowledgements

Data collection and sharing for this project was funded by the Alzheimer’s Disease Neuroimaging Initiative (ADNI) (National Institutes of Health Grant U01 AG024904) and DOD ADNI (Department of Defense award number W81XWH-12-2-0012). ADNI is funded by the National Institute on Aging, the National Institute of Biomedical Imaging and Bioengineering, and through generous contributions from the following: AbbVie, Alzheimer’s Association; Alzheimer’s Drug Discovery Foundation; Araclon Biotech; BioClinica, Inc.; Biogen; Bristol-Myers Squibb Company; CereSpir, Inc.; Cogstate; Eisai Inc.; Elan Pharmaceuticals, Inc.; Eli Lilly and Company; EuroImmun; F. Hoffmann-La Roche Ltd and its affiliated company Genentech, Inc.; Fujirebio; GE Healthcare; IXICO Ltd.; Janssen Alzheimer Immunotherapy Research & Development, LLC.; Johnson & Johnson Pharmaceutical Research & Development LLC.; Lumosity; Lundbeck; Merck & Co., Inc.; Meso Scale Diagnostics, LLC.; NeuroRx Research; Neurotrack Technologies; Novartis Pharmaceuticals Corporation; Pfizer Inc.; Piramal Imaging; Servier; Takeda Pharmaceutical Company; and Transition Therapeutics. The Canadian Institutes of Health Research is providing funds to support ADNI clinical sites in Canada. Private sector contributions are facilitated by the Foundation for the National Institutes of Health (www.fnih.org). The grantee organization is the Northern California Institute for Research and Education, and the study is coordinated by the Alzheimer’s Therapeutic Research Institute at the University of Southern California. ADNI data are disseminated by the Laboratory for Neuro Imaging at the University of Southern California. Data for this study were prepared, archived, and distributed by the National Institute on Aging Alzheimer’s Disease Data Storage Site (NIAGADS) at the University of Pennsylvania (U24-AG041689), funded by the National Institute on Aging. The Alzheimer’s Disease Genetics Consortium (ADGC) supported the collection of samples used in this study through□National Institute on Aging (NIA) grants U01AG032984 and RC2AG036528. Samples from the National Cell Repository for Alzheimer’s Disease (NCRAD), which receives government support under a cooperative agreement grant (U24 AG21886) awarded by the National Institute on Aging (NIA), were used in this study. We thank contributors who collected samples used in this study, as well as patients and their families, whose help and participation made this work possible. The NACC database is funded by NIA/NIH Grant U01 AG016976. NACC data are contributed by the NIA funded ADCs: P30 AG019610 (PI Eric Reiman, MD), P30 AG013846 (PI Neil Kowall, MD), P50 AG008702 (PI Scott Small, MD), P50 AG025688 (PI Allan Levey, MD, PhD), P50 AG047266 (PI Todd Golde, MD, PhD), P30 AG010133 (PI Andrew Saykin, PsyD), P50 AG005146 (PI Marilyn Albert, PhD), P50 AG005134 (PI Bradley Hyman, MD, PhD), P50 AG016574 (PI Ronald Petersen, MD, PhD), P50 AG005138 (PI Mary Sano, PhD), P30 AG008051 (PI Steven Ferris, PhD), P30 AG013854 (PI M. Marsel Mesulam, MD), P30 AG008017 (PI Jeffrey Kaye, MD), P30 AG010161 (PI David Bennett, MD), P50 AG047366 (PI Victor Henderson, MD, MS), P30 AG010129 (PI Charles DeCarli, MD), P50 AG016573 (PI Frank LaFerla, PhD), P50 AG016570 (PI Marie-Francoise Chesselet, MD, PhD), P50 AG005131 (PI Douglas Galasko, MD), P50 AG023501 (PI Bruce Miller, MD), P30 AG035982 (PI Russell Swerdlow, MD), P30 AG028383 (PI Linda Van Eldik, PhD), P30 AG010124 (PI John Trojanowski, MD, PhD), P50 AG005133 (PI Oscar Lopez, MD), P50 AG005142 (PI Helena Chui, MD), P30 AG012300 (PI Roger Rosenberg, MD), P50 AG005136 (PI Thomas Montine, MD, PhD), P50 AG033514 (PI Sanjay Asthana, MD, FRCP), P50 AG005681 (PI John Morris, MD), and P50 AG047270 (PI Stephen Strittmatter, MD, PhD). NIAGADS datasets NG00022, NG00023, and NG00024 contain ADC samples. This paper represents independent research part funded by the National Institute for Health Research (NIHR) Biomedical Research Centre at South London and Maudsley NHS Foundation Trust and King’s College London. The views expressed are those of the author(s) and not necessarily those of the NHS, the NIHR or the Department of Health and Social Care. Funding for genotyping the IRCCS 1 samples was provided by the University of Exeter. IRCCS data was fully supported by “Ministero della Salute”, I.R.C.C.S. Research Program, Ricerca Corrente 2018-2020, Linea n. 2 “Meccanismi genetici, predizione e terapie innovative delle malattie complesse” and by the “5 × 1000” voluntary contribution to the Fondazione I.R.C.C.S. Ospedale “Casa Sollievo della Sofferenza”. The authors are grateful to the investigators of the AddNeuroMed consortium for providing clinical data. AddNeuroMed is funded through the EU FP6 program as part of InnoMed. In addition, we are grateful for additional support from Alzheimer’s Research UK. The authors’ work has been supported in part by the National Institute for Health Research (NIHR) Biomedical Research Centre at South London and Maudsley NHS Foundation Trust and King’s College London. The views expressed are those of the author(s) and not necessarily those of the NHS, the NIHR or the Department of Health. The authors wish to thank the Norwegian registry of persons assessed for cognitive symptoms (NorCog) for providing access to patient data and biological material. NorCog is financed by South-Eastern Norway Regional Health Authority and Norwegian National Advisory Unit on Ageing and Health. The organization and the data collection of HMS cohort has been funded by the Norwegian Institute of Public Health, the Norwegian University of Science and Technology (NTNU), Nord-Trøndelag Hospital Trust and Innlandet Hospital Trust. The REDIC-NH study was administrated by the research centre for Age-related Functional Decline and Disease, Innlandet Hospital Trust, and was initiated by the Norwegian Health Directorate, which also provided funding for the data collection. The data collection of the SAM-AKS cohort has been funded by the research centre for Age-related Functional Decline and Disease, Innlandet Hospital Trust. The authors are grateful to deCODE Genetics for performing the genotyping in the Norwegian cohorts.

